# Quantitative Spatial Analysis of Matrix Remodelling in Wounds Using Digital Pathology and Fractal Analysis

**DOI:** 10.64898/2026.11.04.764252

**Authors:** Xuyang Zhang, Arame Raoofi Arani, Jeremy A. Herrera, Nataliia Marienko, Connor J. Robinson, Svitlana Kurinna

**Author notes:** Correspondence: Svitlana Kurinna.

## Abstract

Wound healing involves spatially heterogeneous changes in matrix composition and organisation that can be difficult to capture using qualitative histological assessment alone. We developed a reproducible digital pathology workflow to quantify matrix features in cutaneous wound sections, compare them with unwounded skin, and derive an exploratory Histological Wound Index (HWI).

Four normal-skin and six wound sections from patients with underlying vascular disease or diabetes were analysed using QuPath, ImageJ/FracLac and Python as a proof of principle. Approximately 500 × 500 µm regions of interest (ROI) were assigned to normal skin, wound centre, proximal edge or distant edge. Collagen, acidic mucins, proteoglycans, glycosaminoglycans (GAGs) and necrotic tissue coverage were quantified using image classifiers and analysed with linear mixed-effects models. Collagen architecture was assessed from collagen-selective binary masks using fractal dimension and lacunarity. Collagen coverage, fractal dimension, mucins coverage, necrosis and lacunarity were z-standardised and integrated into the HWI.

Unwounded skin showed high collagen coverage and fractal dimension, with low lacunarity, mucin, proteoglycans, GAGs, and necrosis. In contrast, wound-centre tissue showed low collagen coverage and fractal dimension together with higher mucin, proteoglycans, GAGs, and necrosis. Distant-edge regions retained comparatively normal-like characteristics, whereas proximal-edge regions showed intermediate values. HWI was highest in normal skin and lowest in the wound centre. A descriptive matched-ROI comparison of two wound sections reproduced this regional ordering when all component measures were derived from identical ROI boundaries.

This workflow provides a reproducible framework for quantifying regional ECM features and collagen-mask architecture in wound histology. The HWI is an exploratory histological summary requiring validation in larger longitudinal datasets.

## Introduction

Wound repair requires coordinated haemostasis, inflammation, proliferation, and remodelling. These phases overlap and are shaped by local cell populations, inflammatory signals, mechanical forces, vascular supply, and extracellular-matrix (ECM) composition (Eming, Martin and Tomic-Canic, 2014). Restoration of tissue integrity requires not only epithelial closure but also an organised dermal environment capable of supporting normal function.

Collagen is a major structural component of skin and contributes to tissue strength, fibroblast migration, and later scar maturation. During repair, collagen is deposited, remodelled and progressively aligned as the provisional matrix matures (Mathew-Steiner, Roy and Sen, 2021). ECM remodelling involves a transition from a provisional, hydrated matrix to a more collagen-rich and mechanically stable dermis; disruption of this process can contribute to abnormal scarring (Xue and Jackson, 2015). However, collagen abundance alone does not fully describe tissue organisation. A collagen-rich region may still contain fragmented or unevenly distributed collagen. Mucins/ground-substance-rich tissue and necrotic material provide additional histological descriptors of altered ECM composition and tissue damage, although they do not identify specific molecules or pathways (Aya and Stern, 2014; Barker and Engler, 2017; Diller and Tabor, 2022; Robinson et al., 2024).

Wound tissue is spatially heterogeneous. The wound centre, wound-adjacent tissue, and more distant peripheral tissue can represent distinct local histological environments. Digital pathology enables these regions to be defined and quantified reproducibly (Robinson et al., 2025), but appropriate analysis must account for the nested relationship between image tiles, ROIs and sections (Bankhead et al., 2017; Riber-Hansen, Vainer and Steiniche, 2012).

Movat’s pentachrome (modified Russell–Movat) simultaneously differentiates five key tissue components within a single section: ground substance (mucin, proteoglycans, and glycosaminoglycans; blue, called mucins for the rest of the paper), elastic fibres and nuclei (black), collagen fibres (yellow), muscle (red), and fibrin (bright red). In chronic skin wounds, this enables comprehensive evaluation of ECM degradation, collagen deposition, elastic fibre fragmentation, proteoglycan-rich granulation tissue, and pericapillary fibrin cuffs.

Fractal dimension and lacunarity provide complementary descriptors of collagen-mask architecture. Fractal dimension describes space-filling complexity, whereas lacunarity describes variation in the distribution of collagen-positive and collagen-negative areas. Such measures have been used in wound and scar studies, although their interpretation depends on imaging modality, specimen preparation and the definition of the analysed collagen signal (Khorasani et al., 2011; Tian et al., 2021; Xu et al., 2022). We developed a reproducible computational pathology workflow to quantify regional ECM composition and collagen-selective mask architecture in normal and wound skin. We hypothesised that normal skin would show the most collagen-rich and structurally organised phenotype, whereas wound-centre tissue would show the greatest disruption of the matrix structure. We also propose an exploratory Histological Wound Index (HWI) that combines the five composition and architecture measures, each oriented so that higher values indicate a more normal-like tissue phenotype.

## Results

ECM coverage analysis included 30 ROI-level observations per tissue component: 12 normal-skin ROIs and six ROIs in each wound-associated region. Fractal analysis included 30 collagen-selective masks. The HWI dataset contained 22 matched section-region observations: four normal skin biopsies (the three ROIs from each normal section averaged to a single value), six wound centre, six proximal edge and six distant edge observations. Three wound-centre masks had a foreground fraction below 1%; they were retained in the primary analysis but interpreted with caution, because lacunarity estimates from very sparse masks are unreliable. A summary of the included sections and ROIs is provided in Table 1. Representative whole-section images and collagen-selective binary masks are shown in Figures 2 and 3, respectively.

**Figure 1.**
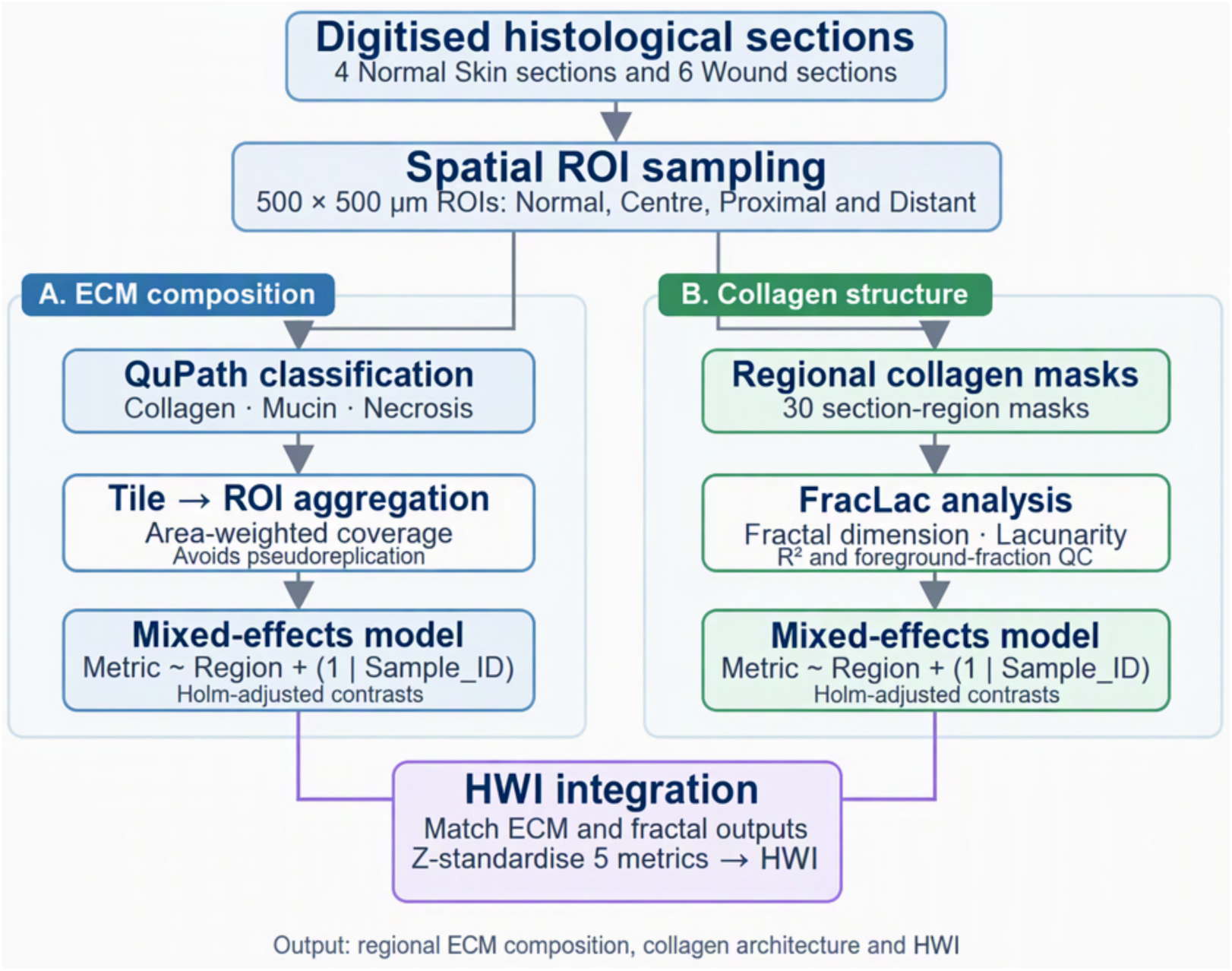
Overview of the analytical workflow. Digitised histological sections were analysed in QuPath to quantify collagen, mucins and necrotic tissue coverage within tiled ROIs. Tile-level values were aggregated to tissue-area-weighted ROI-level measurements and analysed using mixed-effects models. Collagen-selective masks were analysed using FracLac to obtain fractal dimension, lacunarity, R² and foreground fraction. ECM and fractal measurements were matched by Sample_ID and anatomical region, *z*-standardised, and integrated into the HWI.

**Figure 2.**
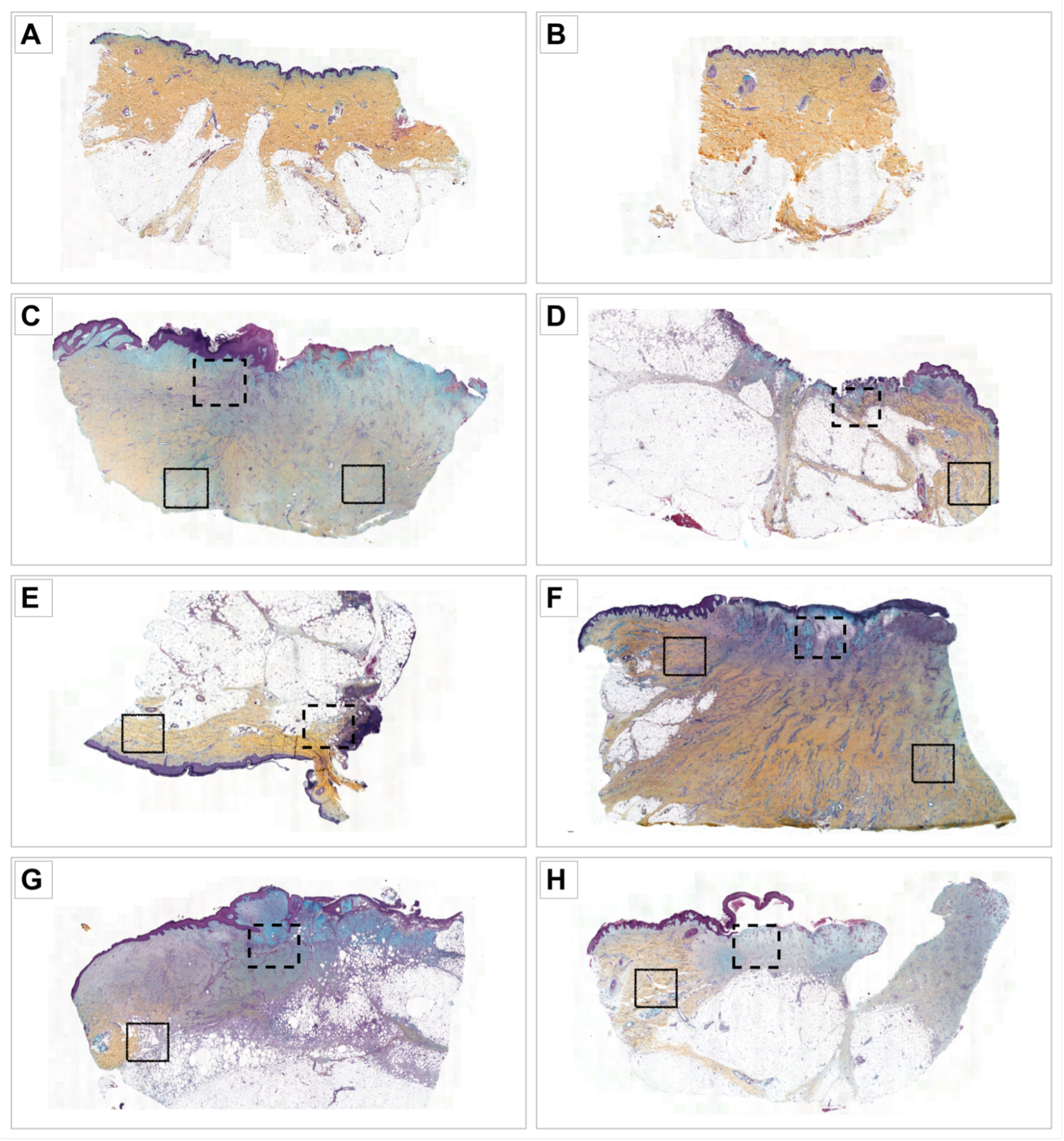
Whole-section pentachrome-stained histological images included in the study. (A–B) Normal skin sections Normal_08 and Normal_09. (C–H) Wound sections Wound_05_1, Wound_05_2, Wound_06, Wound_07, Wound_08 and Wound_09. Collagen-rich connective tissue is yellow/orange, and mucins/ground substance appears blue/turquoise. Necrotic tissue was identified by loss of normal architecture, reduced cellular and nuclear detail, and amorphous non-viable material. Dash-lined boxes show examples of regions central or proximal to the wounded site, while solid-line boxes indicate examples of distant wound edges.

**Figure 3.**
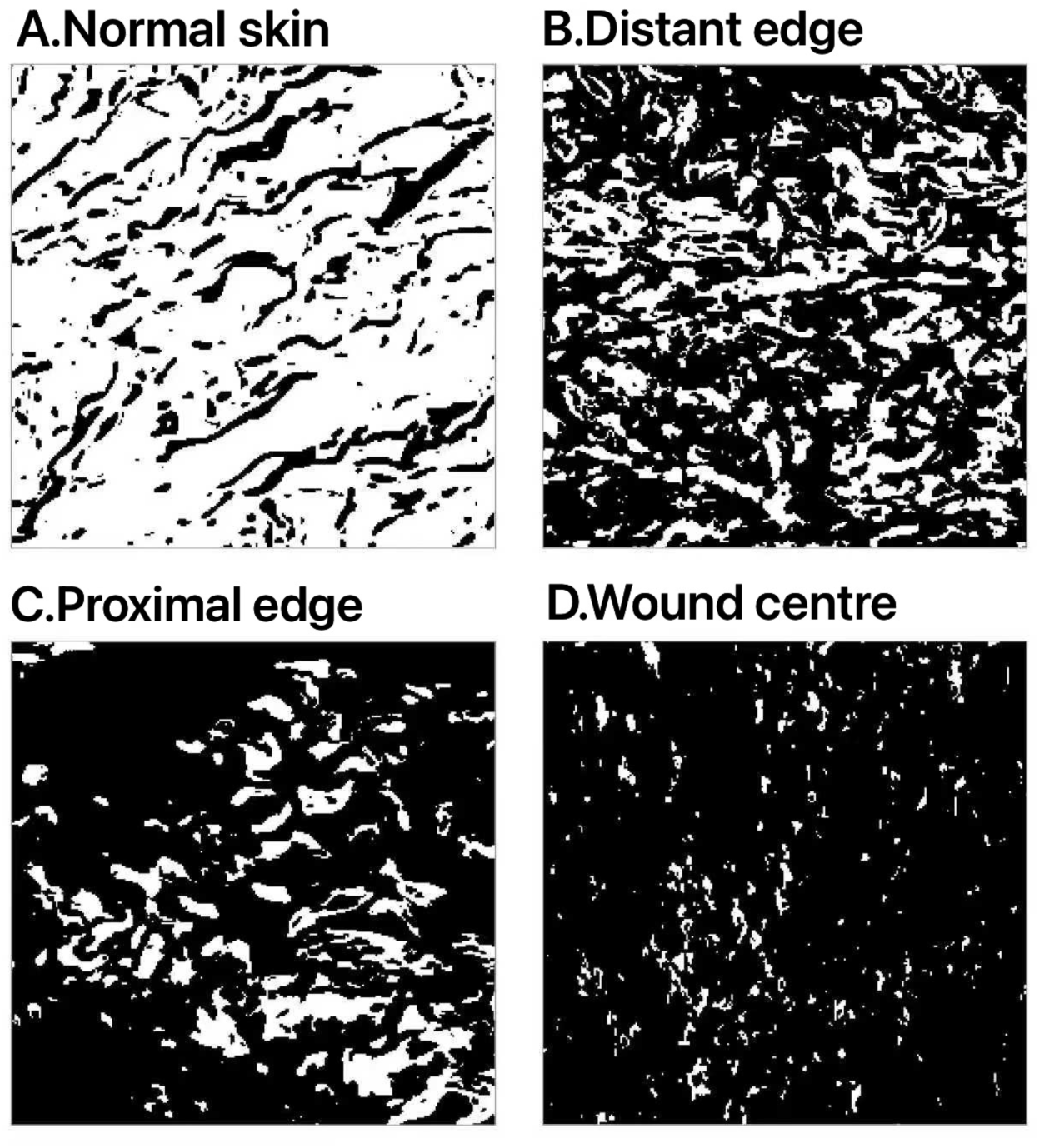
Representative collagen-selective binary masks used for fractal analysis. Representative masks (approximately 500 × 500 µm) from (A) normal skin, (B) distant edge, (C) proximal edge and (D) wound centre are shown. White pixels represent collagen-positive foreground, and black pixels represent background. The masks illustrate regional variation in collagen-positive tissue, with lower coverage and greater spatial disruption in the wound centre than in normal skin. Each mask was analysed using FracLac to derive fractal dimension, lacunarity, R² and foreground fraction.

**Table 1.** Summary of histological sections, anatomical regions, and ROIs included in the study.

| Section group | Section IDs | Number of sections | Anatomical region(s) | ROIs per section | Total ROIs |
|---|---|---|---|---|---|
| **Normal skin** | Normal_06, Normal_07, Normal_08, Normal_09 | 4 | normal skin | 3 | 12 |
| **Wound sections** | Wound_05_1, Wound_05_2, Wound_06, Wound_07, Wound_08, Wound_09 | 6 | wound centre; proximal edge; distant edge | 3<br>(one per region) | 18 |
| **Total** | — | 10 | four analytical regions | — | 30 |

Normal skin had the highest mean collagen coverage (mean ± SD, 61.91 ± 16.89%), followed by distant edge (27.36 ± 16.02%), proximal edge (18.31 ± 15.03%) and wound centre (10.83 ± 13.93%; Figure 4A; Table 2). Compared with normal skin, collagen coverage was lower in the wound centre (Holm-adjusted p = 1.20 × 10⁻⁷), distant edge (p = 2.04 × 10⁻⁴) and proximal edge (p = 5.54 × 10⁻⁶).

**Figure 4.**
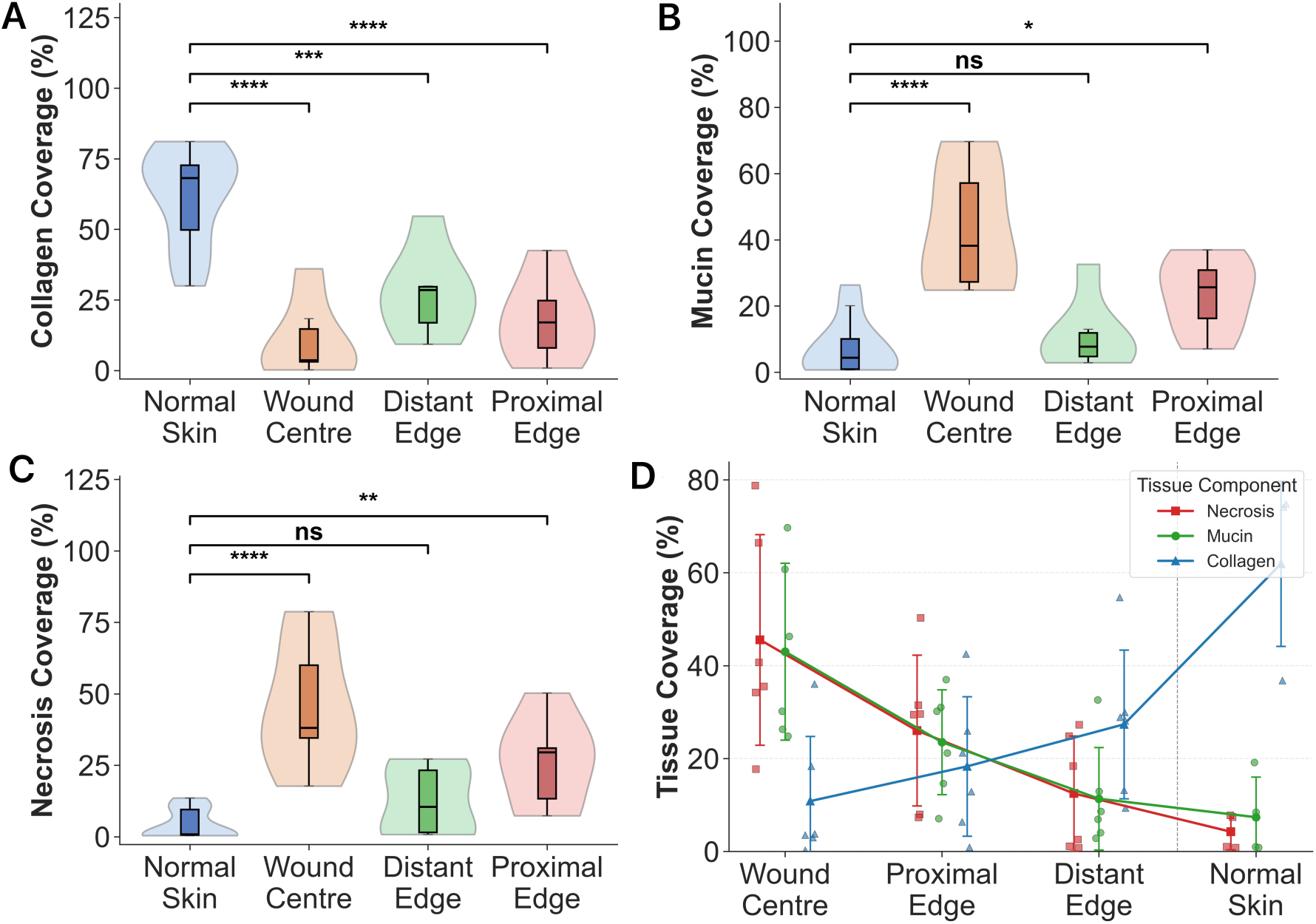
Regional variation in ECM tissue coverage and descriptive ECM profile. (A) Collagen, (B) mucins and (C) necrotic tissue coverage across normal skin, wound centre, distant edge and proximal edge. Violin and box plots show tissue-area-weighted ROI-level observations. Statistical annotations represent Holm-adjusted mixed-effects comparisons with normal skin. (D) Individual section-region values and mean ± standard deviation; lines connect regional means for visual guidance only.

**Table 2.** ROI-level descriptive statistics and Holm-adjusted mixed-effects comparisons for collagen, mucins and necrotic tissue coverage.

| Metric | Region | n | Mean ± SD (%) | Difference to normal skin, % | Holm-adjusted p-value |
|---|---|---|---|---|---|
| Collagen | normal skin | 12 | 61.91 ± 16.89 | — | — |
| Collagen | wound centre | 6 | 10.83 ± 13.93 | −51.08 | 1.20 × 10<sup>−7</sup> |
| Collagen | proximal edge | 6 | 18.31 ± 15.03 | −43.61 | 5.54 × 10<sup>−6</sup> |
| Collagen | distant edge | 6 | 27.36 ± 16.02 | −34.55 | 2.04 × 10<sup>−4</sup> |
| Mucins | normal skin | 12 | 7.37 ± 8.49 | — | — |
| Mucins | wound centre | 6 | 43.01 ± 19.05 | 35.64 | 2.63 × 10<sup>−8</sup> |
| Mucins | proximal edge | 6 | 23.51 ± 11.29 | 16.14 | 1.83 × 10<sup>−2</sup> |
| Mucins | distant edge | 6 | 11.35 ± 11.02 | 3.98 | 0.52 |
| Necrotic tissue | normal skin | 12 | 4.24 ± 5.19 | — | — |
| Necrotic tissue | wound centre | 6 | 45.56 ± 22.66 | 41.32 | 2.60 × 10<sup>−7</sup> |
| Necrotic tissue | proximal edge | 6 | 26.02 ± 16.23 | 21.78 | 9.56 × 10<sup>−3</sup> |
| Necrotic tissue | distant edge | 6 | 12.48 ± 12.40 | 8.24 | 0.29 |

Mucins coverage was lowest in normal skin (7.37 ± 8.49%) and highest in the wound centre (43.01 ± 19.05%). It was higher than normal skin in the wound centre (p = 2.63 × 10⁻⁸) and proximal edge (p = 1.83 × 10⁻²), but not at the distant edge (p = 0.521; Figure 4B; Table 2). Necrotic tissue coverage followed a similar distribution: normal skin 4.24 ± 5.19%, wound centre 45.56 ± 22.66%, proximal edge 26.02 ± 16.23% and distant edge 12.48 ± 12.40%. Necrosis was higher than in normal skin in the wound centre (p = 2.60 × 10⁻⁷) and proximal edge (p = 9.56 × 10⁻³), but not at the distant edge (p = 0.29; Figure 4C; Table 2). Individual section-region values and regional means are shown in Figure 4D.

Wound composition and architectural measures used alongside fractal dimension and lacunarity to characterise regional differences in ECM remodelling could be compared across wound-associated regions and normal skin to assess changes in collagen complexity and overall tissue phenotype. The direction of fractal measures cannot be directly compared across methods because imaging modalities, image dimensionality, preprocessing and the physical signal being measured differ. In this study, fractal dimension and lacunarity are therefore interpreted as two-dimensional descriptors of collagen-selective binary-mask architecture.

Normal skin had the highest mean fractal dimension (1.817 ± 0.026), followed by distant edge (1.726 ± 0.016), proximal edge (1.600 ± 0.091), and wound centre (1.298 ± 0.229; Figure 6A; Table 3). Relative to normal skin, fractal dimension was lower in wound-centre (Holm-adjusted p = 3.31 × 10⁻²⁰) and proximal-edge masks (p = 1.91 × 10⁻⁴), but not significantly different at the distant edge (p = 0.10).

**Figure 5.**
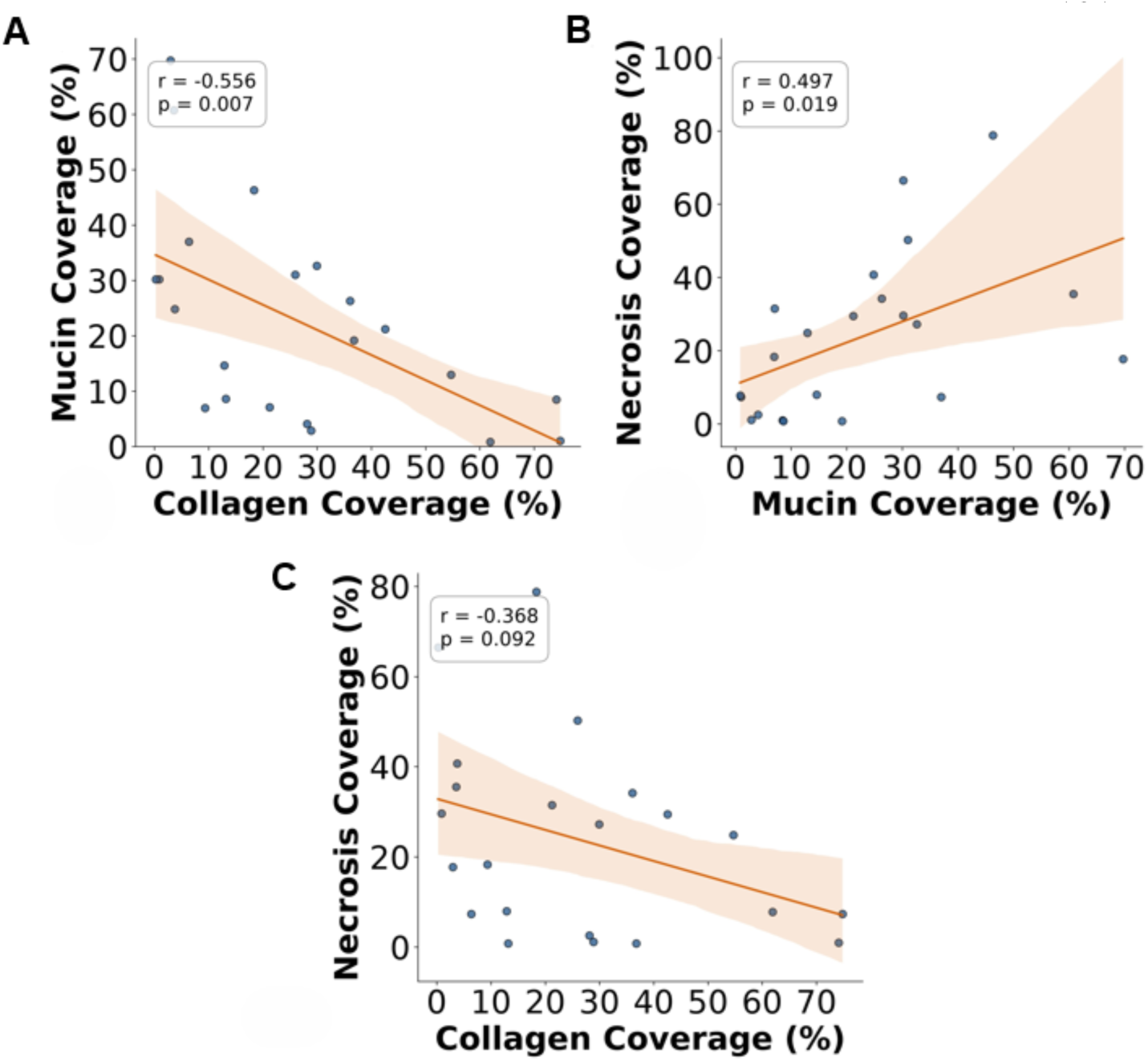
Pairwise correlations between ECM tissue-component coverage. (A) Collagen versus mucins, (B) mucins versus necrotic tissue, and (C) collagen versus necrotic tissue. Each point is one matched section-region observation. Lines are ordinary least-squares fits, and shaded areas are 95% confidence intervals.

**Figure 6.**
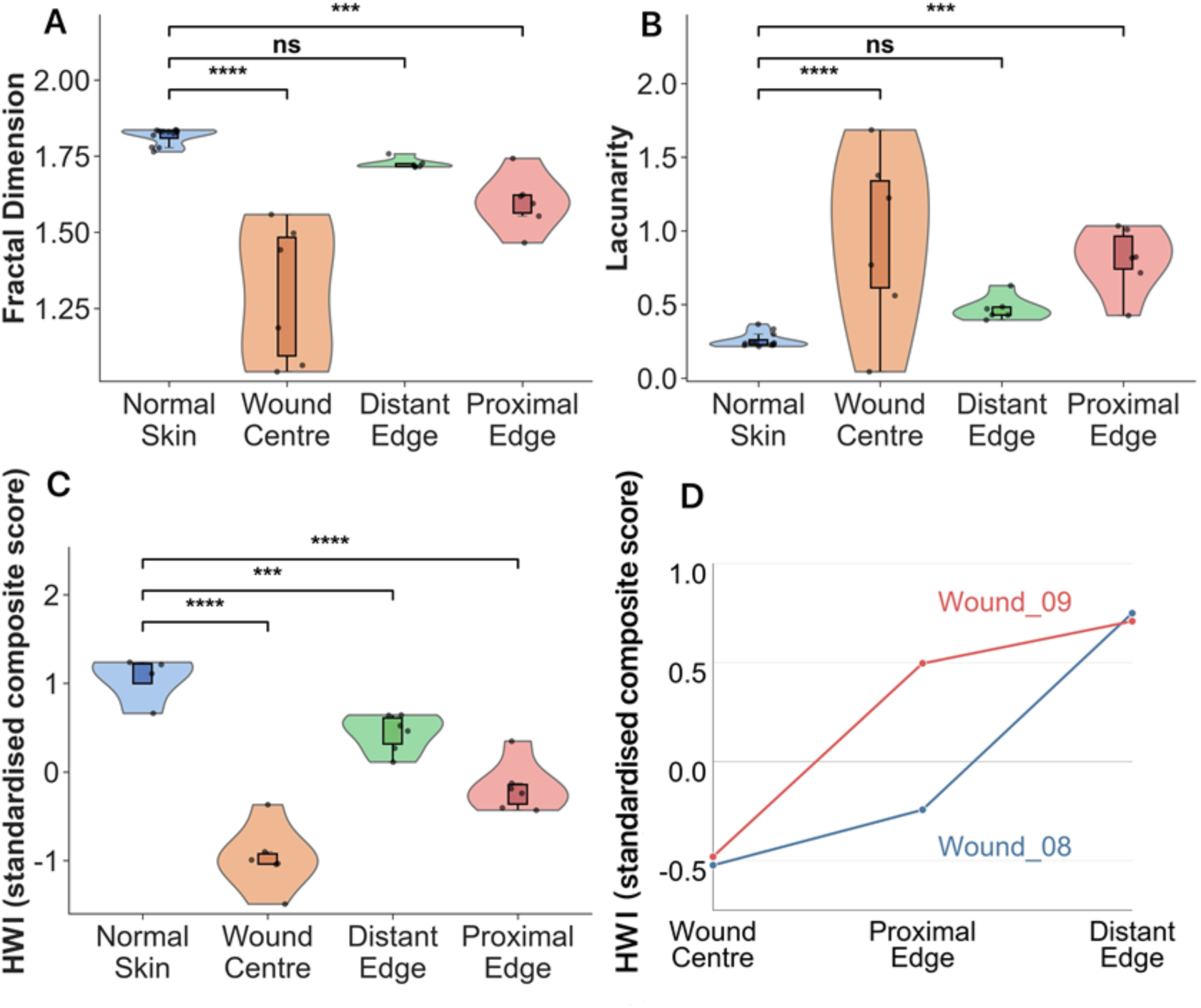
Regional collagen architecture and HWI. (A) Fractal dimension, (B) lacunarity and (C) HWI across normal skin and wound-associated regions. Statistical annotations represent Holm-adjusted mixed-effects comparisons with normal skin. (D) Descriptive co-localised ROI HWI comparison between Wound_08 and Wound_09; lines connect regions within a section and are not fitted models.

**Table 3.** Descriptive statistics and Holm-adjusted mixed-effects comparisons for fractal dimension and lacunarity.

| Metric | Region | n | Mean ± SD | Difference to normal skin | Holm-adjusted p-value |
|---|---|---|---|---|---|
| Fractal dimension | normal skin | 12 | 1.817 ± 0.026 | — | — |
| Fractal dimension | wound centre | 6 | 1.298 ± 0.229 | −0.518 | 3.31 × 10<sup>−20</sup> |
| Fractal dimension | proximal edge | 6 | 1.600 ± 0.091 | −0.217 | 1.91 × 10<sup>−4</sup> |
| Fractal dimension | distant edge | 6 | 1.726 ± 0.016 | −0.091 | 1.01 × 10<sup>−1</sup> |
| Lacunarity | normal skin | 12 | 0.258 ± 0.049 | — | — |
| Lacunarity | wound centre | 6 | 0.944 ± 0.600 | 0.686 | 1.72 × 10<sup>−6</sup> |
| Lacunarity | proximal edge | 6 | 0.805 ± 0.222 | 0.547 | 1.36 × 10<sup>−4</sup> |
| Lacunarity | distant edge | 6 | 0.474 ± 0.083 | 0.216 | 1.16 × 10<sup>−1</sup> |

Lacunarity was lowest in normal skin (0.258 ± 0.049) and highest in the wound centre (0.944 ± 0.600), with intermediate distant-edge (0.474 ± 0.083) and proximal-edge values (0.805 ± 0.222; Figure 6B; Table 3). It was higher than in normal skin in wound-centre (p = 1.72 × 10⁻⁶) and proximal-edge masks (p = 1.36 × 10⁻⁴), but not significantly different at the distant edge (p = 0.12).

HWI was highest in normal skin (1.056 ± 0.268), followed by distant edge (0.441 ± 0.212), proximal edge (−0.174 ± 0.282) and wound centre (−0.971 ± 0.360; Figure 6C). All wound-associated regions differed from normal skin after Holm adjustment.

In the matched-ROI comparison, both Wound_08 and Wound_09 showed the same regional HWI: lowest in the wound centre and highest at the distant edge (Figure 6D). The proximal-edge values differed most between the wounds (Table 4). This two-section analysis was descriptive; no additional inferential testing was performed. The corresponding whole-section images are shown in Figure 7.

**Figure 7.**
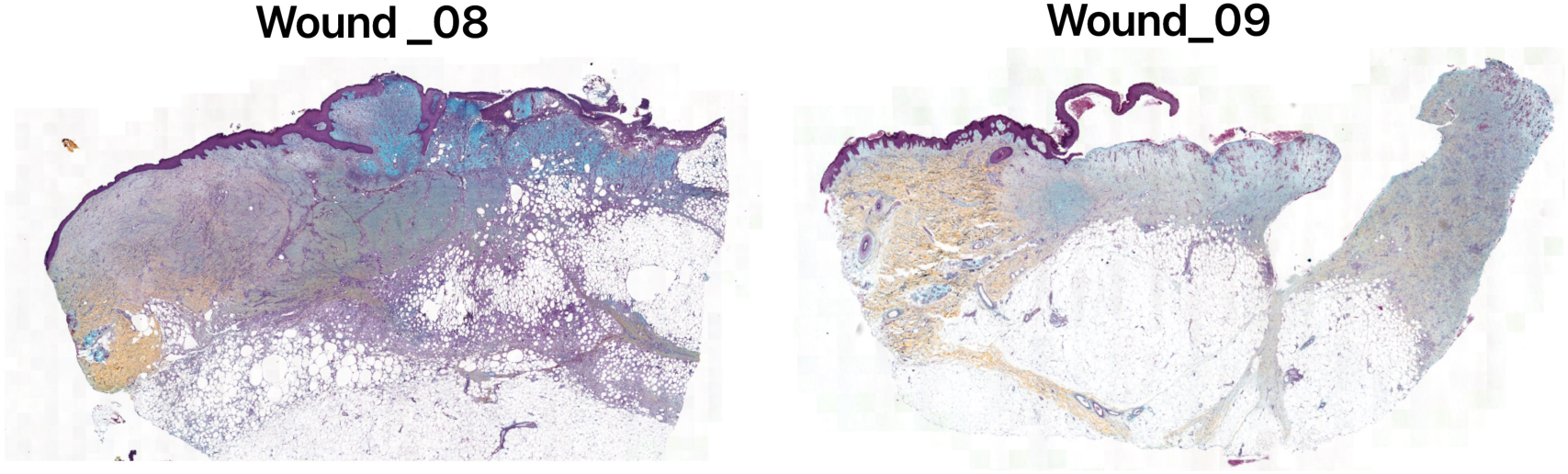
Whole-section pentachrome-stained images of the two wound sections used in the matched-ROI HWI comparison. (A) Wound_08 and (B) Wound_09.

**Table 4.** ECM, fractal and HWI measurements for sampled wounds.

| Sample ID | Region | Collagen (%) | Mucins (%) | Necrosis (%) | Fractal dimension | Lacunarity | Foreground fraction (%) | HWI |
|---|---|---|---|---|---|---|---|---|
| Wound_08 | wound centre | 0.00 | 12.77 | 46.13 | 1.042 | 0.045 | 0.39 | −0.523 |
| Wound_08 | proximal edge | 0.71 | 7.04 | 2.44 | 1.183 | 0.700 | 1.09 | −0.243 |
| Wound_08 | distant edge | 33.09 | 4.62 | 1.65 | 1.749 | 0.352 | 33.30 | 0.749 |
| Wound_09 | wound centre | 0.25 | 26.19 | 7.40 | 1.152 | 0.602 | 0.54 | −0.481 |
| Wound_09 | proximal edge | 24.59 | 7.65 | 1.33 | 1.694 | 0.553 | 24.84 | 0.495 |
| Wound_09 | distant edge | 30.10 | 3.51 | 0.76 | 1.752 | 0.426 | 30.34 | 0.709 |

Collagen coverage was quantified using the QuPath pixel classifier and aggregated using tissue-area weighting within each ROI. Foreground fraction was the percentage of all pixels in the binary mask analysed by FracLac that were white (collagen-positive). The measurements were obtained through separate image-processing workflows and therefore are not numerically identical, even though derived from the same matched ROI (Table 4).

Because pentachrome staining is not routinely used, we also tested whether the same analysis could be applied to consecutive H&E-stained sections. To test this, fractal analysis was repeated on collagen masks from consecutive H&E sections (Figure 8). In unwounded skin, both stains gave nearly identical collagen areas, fractal dimensions and lacunarity (H&E vs pentachrome: collagen 70% vs 68%; D 1.83 vs 1.82; Figure 8 A–D). In wounds, this agreement was lost. H&E masks contained substantially more collagen-positive area than pentachrome masks (49% vs 17%), and most wound ROIs showed a higher fractal dimension and lower lacunarity on H&E (Figure 8 C, D).

**Figure 8.**
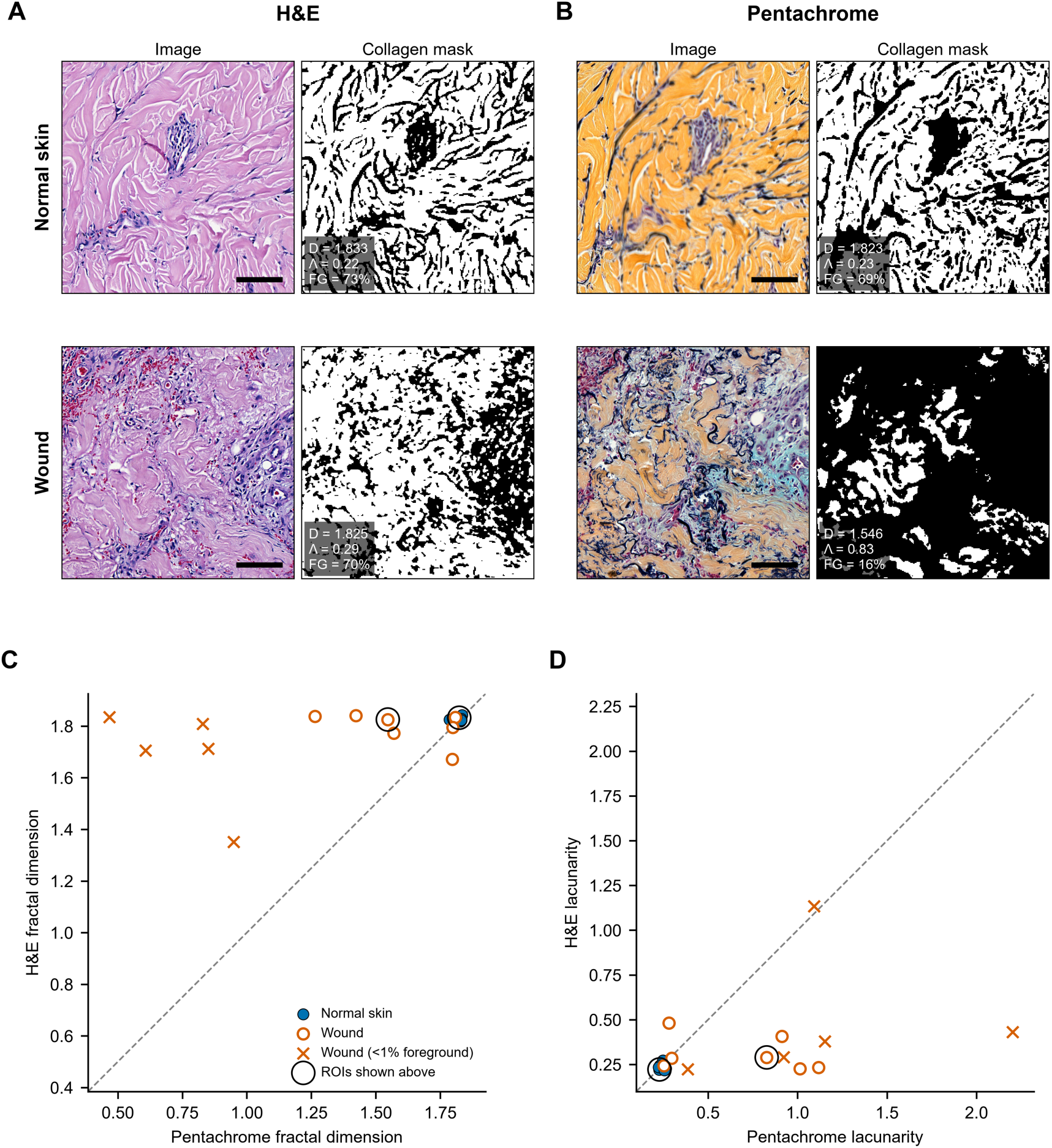
Fractal analysis of H&E corresponds to pentachrome only in unwounded skin. (A, **B)** Representative images and binary collagen masks from consecutive H&E (A) and pentachrome (B) sections of unwounded skin (Normal_06, ROI 1, top) and wound (Wound_05_2, Wound Centre, bottom). Collagen is white in the masks; in pentachrome only yellow-stained (mature) collagen was included. Values on each mask give fractal dimension (D), lacunarity (Λ) and collagen area as a percentage of the ROI (FG). Scale bars, 100 µm. (C, D) Fractal dimension (C) and lacunarity (D) of each ROI measured on H&E versus pentachrome; dashed line, line of identity. Filled circles, unwounded skin (n = 9 ROIs from 3 samples); open circles, wound (n = 12 of 15 ROIs: centre, proximal edge and distant edge from 5 wounds; 3 ROIs with no collagen in the pentachrome mask could not be analysed); crosses, wound ROIs in which either mask contained <1% collagen. Large circles mark the ROIs shown in A and B.

In normal skin, H&E and pentachrome masks showed nearly identical collagen coverage, fractal dimension and lacunarity, suggesting that routinely stained archives could be used to describe dermal collagen architecture in intact tissue. In wound-associated tissue, this agreement was lost. H&E masks contained more collagen-positive area and, in most ROIs, showed higher fractal dimension and lower lacunarity than pentachrome masks. This reflects a limitation of the stain rather than of the fractal method. Eosin labels mature and immature collagen, fibrin and provisional matrix alike, whereas pentachrome separates yellow-stained collagen from mucin-rich and fibrin-rich components. Because these components are abundant in wound matrix, an H&E-derived collagen mask overstates both the amount and the continuity of collagen. Fractal descriptors of wound tissue are therefore only as specific as the mask from which they are derived.

## Discussion

This study developed a computational pathology workflow to describe spatial ECM differences in wound histology. Normal skin had the most collagen-rich and structurally complex collagen-selective masks, whereas wound-centre tissue showed lower collagen coverage and fractal dimension together with higher mucins-positive tissue, necrosis and lacunarity. Distant-edge tissue retained more normal-like characteristics, while proximal-edge tissue showed intermediate values.

The observed reduction in collagen-positive tissue in wound-associated regions is directionally consistent with human chronic-wound work. A reduced dermal collagen abundance and fibril thickness around venous leg ulcers and diabetic foot ulcers was reported relative to intact skin (Sutcliffe et al., 2017). Although their chronic-wound specimens and imaging approach differ from the present material, both studies support the conclusion that wound-associated tissue can show substantial alteration of dermal ECM organisation. The present findings also reinforce the need for spatial analysis: the distant edge was not equivalent to normal skin but differed from the wound centre and proximal edge in several measures. This spatial interpretation is also consistent with observations of changing ECM deposition and histological architecture at the margins of chronic venous ulcers during healing (Herrick et al., 1992).

Fractal measures complemented coverage measurements by describing the spatial organisation of the collagen-selective foreground. (Khorasani et al., 2011) showed that fractal dimension and lacunarity can distinguish scarred from unwounded dermis in a mouse model. Tian et al. (Tian et al., 2021) used a scattering-based approach to quantify collagen organisation, whereas (Xu et al., 2022) reported distinct fractal signatures across different rat dermal injury states.

The distinction between collagen amount and organisation is important. Fractal analysis of fibrotic lung tissue detected spatial changes not captured by collagen content alone (Casey et al., 2024). Although fibrotic lung tissue is not biologically equivalent to cutaneous wound repair, this finding supports the methodological rationale for combining collagen’s quantity and architectural descriptors. Similarly, (de Melo et al., 2026) combined colour-selective histological collagen analysis with fractal dimension in experimentally induced wounds. Together, these studies support the complementary use of image-derived collagen quantity and architecture, while emphasising that direct comparisons require consistent staining, imaging, and preprocessing.

HWI summarised five composition and architecture measures and reproduced their regional ordering. It should be interpreted as an exploratory summary rather than an independent outcome measure, clinical score or surrogate for healing time.

A shareable pipeline, tissue-area-weighted ROI aggregation, and mixed-effects modelling reduced pseudoreplication from image tiles and retained the hierarchical relationship between ROIs and sections. More broadly, reproducibility in digital pathology depends on transparent reporting of tissue selection, classifier performance, ROI definition, and downstream image-processing steps (Bankhead, 2022).

The study is limited by its small number of sections and donors, possible non-independence of normal and wound sections from the same donor, cross-sectional design, unblinded manual ROI selection, classifier-derived mucins and necrosis measures without quantitative validation, absence of stain normalisation, and incomplete co-localisation of primary ECM and fractal ROIs. The sparse collagen foreground in some wound-centre masks also limits lacunarity interpretation. Future work should validate the workflow in larger longitudinal datasets, use fully co-localised ROIs throughout, and relate ECM architecture to fibroblast, immune-cell and endothelial-cell states.

Together, the digital pathology and fractal-analysis workflow identified marked regional differences in ECM composition and collagen-selective mask architecture between normal skin and wound-associated tissue. The results support the use of pentachrome staining for wound architecture and provide a basis for the analysis of image-derived ECM characteristics in larger datasets.

## Methods

### Histological material and spatial regions

The dataset comprised ten independent digitised pentachrome-stained human skin sections derived from five image files. One file contained two wound sections (Wound_05_1 and Wound_05_2); the remaining files each contained one normal-skin and one wound-skin section. The resulting dataset contained four normal skin and six wound skin sections. All sections were digitised on 30 March 2022. This was a secondary computational analysis of pre-existing digital histology images; no new tissue collection, participant recruitment or intervention was undertaken.

Three spatially separated ROIs (approximately 500 × 500 µm each) were selected from each normal skin section. One approximately 500 × 500 µm ROI was selected from each wound section at the wound centre, proximal edge and distant edge. Wound centre referred to tissue within the internal wound-associated area; proximal edge to tissue immediately peripheral to the wound centre; and distant edge to tissue at a greater relative separation within the same section. Categories were relative to anatomical locations rather than fixed distance-based compartments. ROIs were selected after whole-slide review and avoided folds, tears, large empty areas and other artefacts. Representative whole-section images are provided in Figure 2.

### Digital pathology and ROI aggregation

Sections were analysed in QuPath version 0.6.0 (Bankhead et al., 2017). Three pixel classifiers were developed for this study using manually annotated training regions and the Random Trees algorithm. The Pentachrome_Collagen_v1 classifier identified the yellow/orange collagen-stained signal and distinguished collagen-positive pixels from non-collagen tissue, with white background excluded. The mucins classifier identified the blue/turquoise mucins-positive signal and distinguished mucins-positive pixels from the background. The necrotic tissue classifier was initially trained using colour features to distinguish necrotic tissue from the background. Because necrosis was not represented by a single staining colour, its outputs were additionally reviewed in relation to tissue morphology, including loss of normal tissue architecture, reduced cellular and nuclear detail, and amorphous non-viable material. The classifiers quantified the percentage coverage of collagen, mucins, and necrotic tissue within nominal 100 × 100 µm image tiles. Coverage values were obtained from the QuPath measurement columns Pentachrome_Collagen_v1: collagen %, mucins classifier: mucins %, and necrotic tissue: Necrotic tissue %. Classifier outputs were visually reviewed against the corresponding Pentachrome-stained histology before export to identify background classification, staining artefacts, tissue folds, and cross-classification errors.

Tile-level measurements were aggregated to tissue-area-weighted ROI-level coverage values. This prevented tiles with little analysable tissue from contributing equally to tiles with greater tissue area and avoided treating nested technical tiles as independent biological observations. The overall analytical workflow is summarised in Figure 1.

### Collagen-selective masks and fractal analysis

Thirty collagen-selective binary masks were analysed: 12 from normal skin and 18 from wound-associated regions. The yellow/orange collagen-stained signal was extracted from pentachrome-stained image regions in QuPath, converted into black-and-white masks in ImageJ version 1.54g, and analysed using FracLac version 2015Sep090313a9330 (Karperien, 2015; Schneider, Rasband and Eliceiri, 2012). White pixels represented extracted collagen-stained foreground, and black pixels represented background. FracLac outputs included mean fractal dimension, lacunarity, coefficient of determination (R²) and foreground fraction.

The primary ECM and fractal analyses were matched at Sample_ID × Region level because their ROI boundaries were not necessarily identical. Masks with a foreground fraction below 1% were retained because very low collagen coverage is biologically informative; however, lacunarity estimates from these masks are unstable, so they were flagged and examined in a sensitivity analysis. Representative collagen-selective binary masks are shown in Figure 3.

### Comparison with H&E-stained sections

To test whether collagen architecture could be assessed on routinely stained tissue, a separate set of ROIs was analysed on pentachrome sections and on consecutive H&E-stained sections. These ROIs were selected independently of those used in the primary analysis, using the same region categories, so pentachrome values in this comparison are not identical to those reported above. No H&E section was available for sample 07, which was excluded, leaving 24 paired ROIs of approximately 500 × 500 µm (9 normal skin and 15 wound-associated). Wound ROIs were placed using anatomical criteria without reference to the collagen masks: wound centre midway between the epidermal margins beneath the scab, proximal edge at the wound margin, and distant edge outside the margin, all away from follicles, glands and large vessels. H&E images were registered to the corresponding pentachrome images in QuPath and ROIs were transferred between the registered images, with pentachrome as the reference. Masks from the non-reference stain were resampled into the reference frame using nearest-neighbour interpolation. Collagen was segmented with Pentachrome_Collagen_v1 (yellow-stained collagen only) and a separate Random Trees classifier for H&E (HE_Collagen_v1, eosinophilic dermal collagen), both operating at 1.94 µm/pixel; masks were exported at 0.243 µm/pixel. Masks from both stains were inverted before analysis so that collagen was treated as foreground and analysed in FracLac using box counting on binary images without filtering, 12 grid positions, square boxes and 100 box sizes from 5 pixels to 45% of image width. Agreement between stains was assessed descriptively at ROI level by plotting paired values against the line of identity, separately for normal skin and wound; no inferential testing was performed.

### Statistical analysis and HWI

Collagen, mucins and necrotic tissue coverage were analysed separately using linear mixed-effects models with anatomical region as a fixed effect and Sample_ID as a random intercept. Fractal dimension, lacunarity and HWI were analysed using the same model structure. All models were fitted by maximum likelihood. Planned comparisons contrasted normal skin with wound centre, distant edge and proximal edge. Fixed-effect contrasts were tested using two-sided Wald z-tests based on an asymptotic normal approximation, and Holm adjustment was applied separately within each metric across the three comparisons with normal skin. Random-effect variances and residual diagnostics are reported in Supplementary Table 1. Pearson correlations between section-region summaries of collagen, mucins and necrosis were exploratory.

HWI combined *z*-standardised collagen coverage and fractal dimension as positive components, and mucins coverage, necrotic tissue coverage and lacunarity as negative components. Each component was standardised as:

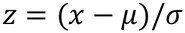

Where x is the observed value, μ is the mean across the matched dataset, and σ is the corresponding standard deviation. HWI was calculated as:

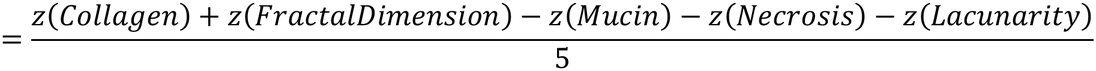

Equal weights were used.

HWI is an exploratory image-derived summary and not a validated clinical or prognostic score. A focused descriptive comparison used one fully co-localised ROI in each of the wound centre, proximal edge and distant edge of Wound_08 and Wound_09. Its z-scores used the reference mean and population standard deviation from the primary 22-observation matched dataset.

### Software and reproducibility

Downstream data processing, statistical analysis and figure generation were performed in Python 3.11 (64-bit) using NumPy (version 1.24 or later), pandas (version 2.0 or later), matplotlib (version 3.7 or later), seaborn (version 0.12 or later), SciPy (version 1.10 or later), statsmodels (version 0.14 or later) and openpyxl (version 3.1 or later). The code, folder structure and dependency instructions are available at https://github.com/595230329/RP2-wound-ECM-fractal-analysis

### Ethics statement

The use of human tissue samples was approved by the NHS Health Research Authority (Greater Manchester Central Research Ethics Committee) through the ComplexWounds@Manchester biobank with informed patient consent under ethical permissions REC23/NW/0320.

### Data and code availability

Analysis code, dependency information and the required input-folder structure are available at https://github.com/595230329/RP2-wound-ECM-fractal-analysis.

## Funding

S.K. is supported by a CDA MRC MR/P009700/1 and MRC TSF MR/X023397/1.

### Author contributions

**Xuyang Zhang:** conducted research, developed the methodology and conceptualized the histological wound index, analyzed data and wrote the manuscript. **Arame Raoofi Arani:** conducted research, contributed to the methodology, analyzed the data and contributed to the writing and editing of the manuscript. **Nataliia Marienko:** designed and assisted the fractal analysis, reviewed the manuscript. **Jeremy Herrera:** experimentally supported wound sections acquisition and staining. **Connor Robinson:** supervised part of the project and conceptualized the software; **Svitlana Kurinna:** conceptualized and supervised the project, acquired funding.

## Acknowledgements

We would like to thank Drs. Jason Wong and Adam Reid for proving the access the paraffin sections from human wound biopsies processed at ComplexWounds@Manchester biobank.

We would like to give special thanks to Ibrar Yunus, Agentic Full-Stack AI Engineer, for reviewing the codes on GitHub repository and helping to improve them.

