## Supplement for "Quantitative Spatial Analysis of Matrix Remodelling in Wounds Using Digital Pathology and Fractal Analysis": Supplementary Information.pdf

Supplementary Table 1. Mixed-effects model variance components and residual diagnostics.

This table summarises the variance components, residual diagnostics and convergence status of the six mixed-effects models for collagen coverage, mucin coverage, necrotic tissue coverage, fractal dimension, lacunarity and HWI. It reports observation and sample-group counts, random-intercept and residual variances, intraclass correlation coefficients (ICC), Shapiro-Wilk statistics, maximum absolute standardised residuals and convergence status. These measures characterise between-sample variation and residual behaviour, helping assess potential departures from model assumptions and interpret the regional comparisons reported in the main text.

| **Outcome** | **n** | **Sections** | **Random-intercept variance** | **Residual variance** | **ICC** | **Shapiro–Wilk W** | **Shapiro–Wilk p-value** | **Maximum \|standardised residual\|** | **Converged** |
|---|---|---|---|---|---|---|---|---|---|
| Collagen coverage | 30 | 10 | 193.344 | 23.956 | 0.890 | 0.973 | 0.619 | 1.850 | Yes |
| Mucin coverage | 30 | 10 | 37.707 | 90.657 | 0.294 | 0.933 | 0.059 | 2.623 | Yes |
| Necrotic tissue coverage | 30 | 10 | 110.032 | 54.949 | 0.667 | 0.975 | 0.673 | 1.703 | Yes |
| Fractal dimension | 30 | 10 | 0.0029 | 0.0075 | 0.283 | 0.934 | 0.065 | 2.315 | Yes |
| Lacunarity | 30 | 10 | 0.0077 | 0.0626 | 0.109 | 0.844 | <0.001 | 3.239 | Yes |
| HWI | 22 | 10 | 0.0167 | 0.0505 | 0.249 | 0.992 | >0.999 | 2.061 | Yes |

Each model included anatomical region as a fixed effect and Sample_ID as a random intercept and was fitted by maximum likelihood. n is the number of section–region/ROI observations entering each model; Sections is the number of independent Sample_ID groups. ICC = random-intercept variance / (random-intercept variance + residual variance).

*Residual diagnostics.* Diagnostics are based on conditional residuals (observed minus fitted values including estimated random effects). Standardised residuals were divided by the residual standard deviation. Shapiro–Wilk results are descriptive because observations are clustered and the sample sizes are small.

*Inference.* Planned fixed-effect contrasts used two-sided Wald z-tests based on an asymptotic normal approximation. Holm adjustment was applied separately within each outcome across the three comparisons with normal skin. Models were first fitted using L-BFGS, with Powell optimisation used as the prespecified fallback when required. Variance components are reported in squared outcome units.
